# Habitat loss-induced tipping points in metapopulations with facilitation

**DOI:** 10.1101/481176

**Authors:** Josep Sardanyés, Jordi Piñero, Ricard Solé

## Abstract

Habitat loss is known to pervade extinction thresholds in metapopulations. Such thresholds result from a loss of stability that can eventually lead to collapse. Several models have been developed to understand the nature of these transitions and how are they affected by the locality of interactions, fluctuations, or external drivers. Most models consider the impact of grazing or aridity as a control parameter that can trigger sudden shifts, once critical values are reached. Others explore instead the role played by habitat loss and fragmentation. Here we consider a minimal model incorporating facilitation along with habitat destruction, with the aim of understanding how local cooperation and habitat loss interact with each other. An explicit mathematical model is derived, along with a spatially explicit simulation model. It is found that a catastrophic shift is expected for increasing levels of habitat loss, but the breakpoint dynamics becomes continuous when dispersal is local. Under these conditions, spatial patchiness is found and the qualitative change from discontinuous to continuous results from a universal behaviour found in a broad class of nonlinear ecological systems (Weissmann and Shnerb, 2014; Martin *et al.* PNAS (2015) E1828-E1836). Our results suggest that species exhibiting facilitation and displaying short-range dispersal will be markedly more capable of dealing with habitat destruction, also avoiding catastrophic tipping points.

## INTRODUCTION

A major threat to the viability of many extant ecosystems is connected to their susceptibility to habitat loss. Among the potential problems associated to this process, a great concern exists in relation to the tempo of ecosystems collapse. A growing consensus among scientists is that species diversity might face the presence of the so-called *catastrophic shifts*, namely discontinuous transitions from a diverse to a poor (or fully extinct) community once parametric thresholds are crossed (Scheffer *et al.* 2001, Scheffer 2009, Rockström *et al.* 2009, Solé 2011). In this context, prediction of future scenarios becomes an issue, given the ongoing degradation of habitats associated to climate change, demographic pressures and the potentially irreversible nature of tipping points (Scheffer *et al.* 2001, Barnosky *et al.* 2012, Barnosky & Hadly 2016).

Mathematical and computational models of ecosystem responses against different classes of damage have been developed in the last decades. Some of these models concern the impact of habitat loss and fragmentation, indicating the existence of destruction levels leading to species extinctions (Bascompte & Solé 1996, Hanski 1999, Solé *et al.* 2002). Others incorporate the impact of increasing aridity levels, which damages soil crusts and reduce the quality of soil and vegetation resilience (Maestre *et al.* 2015). The effective impact level of these two classes of perturbation is tied to the nonlinearities associated to individual interactions. In standard models of habitat destruction, vegetation reacts as a logistic-like growth shape which, combined with linear decay, creates the conditions for extinction thresholds (Levins 1969).

On the other hand, when individuals interact in non-linear ways, responses to increasing levels of aridity or grazing also lead to extinction thresholds, although the nature and speed of the decay towards extinction is very different. A specially relevant example involves the future of semi-arid ecosystems (Rietkerk & van de Koppel 1997, Scanlon *et al.* 2007, Kéfi *et al.* 2007a-b, Solé 2007) where warming, steady declines in rainfall and increased grazing are likely to promote a sudden shift to a desert state (Foley *et al.* 2003). The analysis and modelling of spatial patterning in semiarid habitats consistently supports the suggestion that rapid shifts might occur in a near future (Kéfi *et al.* 2007a-b, Solé 2011).

Several types of models have been proposed to explain ecosystem responses under different environmental stresses. When dealing with drylands, a key factor appears to be the presence of *facilitation processes*, i.e. positive pairwise interactions between individuals leading to the benefit of at least one of the interacting partners (Brooker *et al.* 2008). A specially relevant example involves positive interactions where one organism makes the local environment more favourable for another (which can belong or not to the same species). This positive effect can occur either directly or indirectly. An example of the former includes shading mechanisms that reduce water or nutrient stress. The later would include removing competitors or deterring predators (see Bruno *et al.* 2003). Since both facilitation and habitat loss can occur together and they have distinct types of dynamics: What is the impact of habitat destruction on an ecological system involving facilitation? How does this effect interact with the nonlinearities associated to facilitation? What kind of tipping points (i.e., catastrophic or smooth) are found under habitat loss and facilitation?

Here we explore this problem by presenting a minimal model that captures both components, as well as the spatially extended counterpart under different dispersal regimes and facilitation processes. The mean-field (well-mixed) version is built from a microscopic description based on the underlying rules of facilitation, colonisation and extinction. It predicts that first-order (catastrophic) transitions will occur for increasing levels of habitat loss. However, the spatially explicit versions of the model reveal a rather interesting phenomenon: the transition becomes continuous when interactions are local, confined to nearest neighbours, becoming catastrophic after a given dispersal range is exceeded. This seems to be a rather generic phenomenon, associated to the universal properties of complex systems exhibiting phase transitions (Weissmann & Shnerb, 2014; Villa Martin *et al.* 2014). A detailed analysis of the effects of dispersal is presented and the potential implications for ecosystems’ fate are discussed.

## MATERIALS AND METHODS

### Master equation and mean field model

Our goal is to develop a spatial generalisation of metapopulations with facilitation under habitat destruction. The system introduced here aims to model the colonisation and extinction dynamics of sessile organisms, particularly within the context of drylands and rangelands. Here we consider an individual-based description where interactions are considered in an explicit way. From that microscopic description, average population equations are obtained. We begin by building a formal lattice model using a *master equation* considering the processes of facilitation, colonisation, and decay of the individuals (Fig. 1). Such a lattice model has been used in different contexts (Chopard and Droz, 1998) including field theoretic models of biogeography (O’Dwyer *et al.* 2010; Azaele *et al.* 2016; Pigolotti *et al.* 2018). Let us consider a given habitat described as a two-dimensional grid of size *L*^2^. Three discrete state variables are defined per site: *A*_*i*_, *E*_*i*_, *D*_*i*_ ∈ {0, 1} with *i ∈* {1, *…, L*}^2^. These local variables correspond to the *occupied, empty*, and *destroyed* states of a site, respectively. For example, a site *i* is said to be occupied at a given time *t* if *A*_*i*_ = 1 and *E*_*i*_ = *D*_*i*_ = 0. In order to avoid degeneracy, a state constrain is imposed: *A*_*i*_ + *E*_*i*_ + *D*_*i*_ = 1 for all sites. The following reaction rules determine the dynamics of the system:

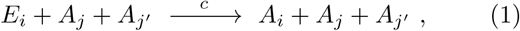

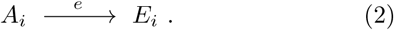

**FIG. 1.**
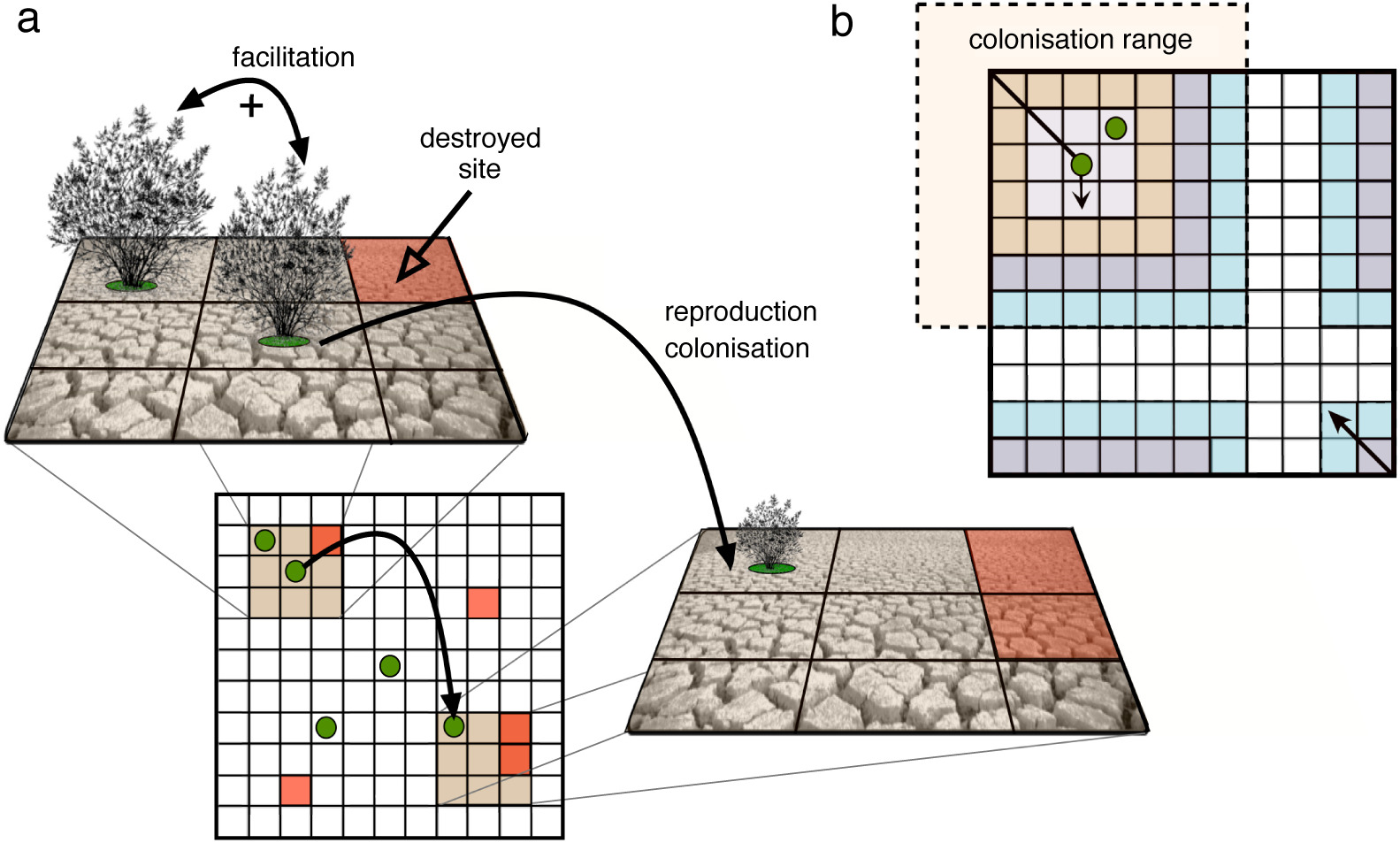
Schematic diagram of the ecological system with facilitation in reproduction under habitat destruction illustrated with the cellular automaton (CA) model. The individuals can grow and reproduce due to the cooperation with neighbouring individuals. (a) The CA is a square lattice and has three states: (i) sites occupied by an individual (green circle), (ii) empty sites that can be colonised (white cells), and (iii) destroyed sites (red cells) where no establishment or growth of an individual is possible. The dynamics of reproduction involves facilitation i.e., an individual can reproduce if a neighbouring place is occupied. Once an individual reproduces, can colonise another empty site of the lattice. If the colonised site is destroyed the individual can not grow. The lattice is assumed to have toroidal boundary conditions. In (b) we display two possible colonisation scenarios: local (short thin arrow) and long-range (solid arrow) dispersal.

Here, reaction (1) denotes the process of colonisation: a site with an occupied state *A*_*j*_ = 1 interacts with an occupied neighbouring site *A*_*j*′_ and colonises and empty site *E*_*i*_ at a probability rate *c*. Hereafter, we will use the notation {*j, j*′}_*i*_ to refer to immediate next neighbours of site *i*. Reaction (2) corresponds to the extinction process,where an occupied site *A*_*i*_ becomes empty at a rate *e*.

For simplicity, let us consider a compact notation by introducing local vector states

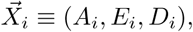

a state vector for global configurations.

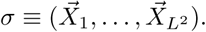

Now, it is possible to define the ensamble of time-evolving configuration probabilities characterised by a distribution function *P* (*σ*; *t*). Considering reactions (1)-(2), it can be shown that the probability distribution evolves under the following master equation (see Section S3.1). This equation is described, in general terms, as follows (van Kampen 1981, Méndez *et al.* 2014):

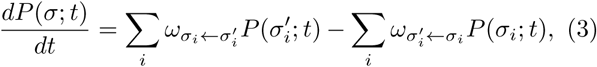

where the normalisation condition Σ _*σ*_ (*σ*; *t*) = 1, holds. Here the whole set of transitions between different probabilistic configurations are taken into account.

The first term at the right hand side of Eq. (3) includes all the favourable transitions towards *P* (*σ*; *t*) whereas the second term stands for all changes going out from it. In our mathematical framework, the state space incorporates the lattice description.

For the specific set of rules given by reactions (1)-(2), it can be shown (see Section S3.1) that the resulting master equation is given by

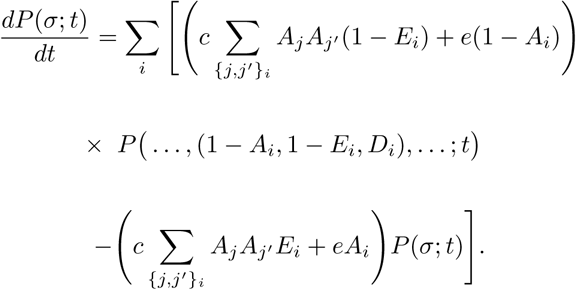

Averages of any (local or global) observable magnitude, namely 𝒪, may now be computed as 〈𝒪〉 (*t*) = Σ _*σ*_ 𝒪 *P* (*σ*; *t*). Furthermore, the time-derivatives of a macroscopical observable 〈𝒪〉 (*t*) can be derived from Eq. (3). Thus, for the average activity of the system we obtain

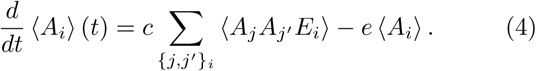

Finally, we perform first order calculations on Eq. (4) by ignoring all spatial correlations. This process will lead us to a so-called mean field theory, upon which a simple analysis becomes feasible. Recalling the non-degeneracy constain *E*_*i*_ = 1– *D*_*i*_ *–A*_*i*_, and breaking correlations by assuming that the three-point functions may be approximated as 〈𝒪_*i*_ *𝒪*_*j*_ *𝒪*_*k*_〉 ∼ *〈𝒪 〉*_*I*_,〈𝒪_*j*_ *〈,⟨𝒪*_*k*_, 〉 then Eq. (4) turns into the following ordinary differential equation

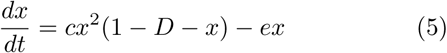

where we have introduced the following notation: 〈*A*_*i*_ 〉= *x*, and 〈*D*_*i*_ 〉= *D*. Effectively, by this procedure we are mixing the system, or, alternatively, connecting all the sites together. Hence, Eq. (5) does not reflect any intrinsic spatial effects. The last equation can be considered as a modified version of Levins’ metapopulations model with habitat destruction (see Section S1). The nonlinear term on population growth and colonisation for this model is interpreted as a process of autocatalysis, where the reproduction kinetics (given by *cx*^2^) is not exponential but hyperbolic. When no habitat fragmentation is considered, i.e. *D* = 0, then it becomes equivalent to an autocatalytic replicator with exponential degradation (see Windus & Jensen 2007, Sardanyés & Solé 2007, Fontich & Sardanyés 2008).

### Spatial stochastic computational model

Following computational methods of stochastic spatial systems we will implement the dynamics tied to reactions (1) and (2) using a cellular automaton (CA). The CA is given by a square *L×L* lattice of side size *L* and periodic boundary conditions (see Fig.1). The states of the CA are given by sites that can be occupied (*S*_*a*_); empty (*S*_*e*_); or destroyed (*S*_*d*_). The fraction of destroyed sites (*D*) is implemented by randomly distributing the given number of *S*_*d*_ states over the grid during the lattice initialisation (see below). The states *S*_*d*_ do not follow any dynamics, i.e., once the system is initialised, only the non-destroyed sites evolve dynamically.

The CA algorithm works as follows: at each time generation, *τ, L ×L* random sites are chosen and a stochastic version of the state-transitions rules following reactions (1) and (2) is applied. This updating process ensures that, on average, all cells in the lattice will be updated once every generation. The state-transition rules are:

- *Reproduction and colonisation*: a site, say *S*_*i*_, is randomly chosen. If *S*_*i*_ is empty or destroyed nothing happens. If *S*_*i*_ contains a species (is occupied), a random nearest neighbour (using a Moore neighbourhood) is selected, say *S*_*j* *≠i*_. If *S*_*j* *≠i*_ contains another species, the species in *S*_*i*_ reproduces and colonises another randomly chosen neighbour site *S*_*k≠i,j*_, placed within a given colonisation distance, *δ* (see below). Colonisation occurs with probability *c∈* [0, 1]. If *S*_*k* *≠i,j*_ is a destroyed or an occupied site, no colonisation takes place.
- *Extinction*: a random cell is chosen. If the cell is occupied by a species, it dies with probability *e* ∈ [0, 1].

The initial configuration of the CA states is given by a random distribution of empty and occupied states, if not otherwise specified. Then, a fraction of destroyed sites (*D*) is introduced randomly over the lattice.

The previous rules consider that the process of reproduction and further colonisation of a given individual (occupied site) occurs whenever it is surrounded by other individuals that facilitate this process (Fig. 1). Alternatively, it is possible to postulate a different type of facilitation process by considering that facilitation takes place after colonisation, i.e., supposing that seeds establishment is limited by the presence of neighbours around the colonised site. For this case, reproduction is not density-dependent. We briefly discuss this alternative model in the last part of the Results Section. A full description of this model can be found in Section S4 (see also Fig. S7).

As mentioned above, colonisation is implemented at a given dispersal distance, varying from local (closest neighbours) to long-range (any lattice site) ranges. The former would be more limited to shorter dispersal, while the latter may colonise over longer distances. In order to consider the full spectrum between these two extreme we use the approach by Wodarz & Levy (2011). Here, authors used a function to obtain the distance over which colonisation can take place. This distance is determined by

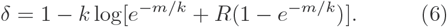

This function provides a random number between 1 and *m*, where *m* is the maximum distance that can be achieved during colonisation. Since the CA model has periodic boundary conditions we shall assume that *m* = *L/*2. *R* is a uniformly distributed random number, and the parameter *k* determines how steep the distribution of the resulting random number *δ* is (see Fig. S2). If *k →* 0, the distribution of *D* is very steep and the probability of colonisation declines very fast with distance. When *k* = 0, *δ* = 1 and thus colonisation occurs within the nearest neighbours (local range colonisation). On the other hand, if *k → ∞*, then the distribution of *δ* tends to be uniform and any position in the lattice has the equal chance of being colonised (long-range colonisation). The latter corresponds to the perfect mixing extreme which is equivalent to the mean field model given by Eq. (5). Intermediate values of *k* allow for a continuum between these two extremes.

In the spatial simulations developed in this work, *k* will be used as a control parameter to test how the nature of the transition towards the extinction of the metapopulation depends on the colonisation ranges. We notice that the distance *δ* in our system will be implemented as a Chebyshev distance (see Fig. 1b). Since *k* does not provide clear information about the colonisation distance, we use mean colonisation distance for any given value of *k*, namely 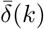, as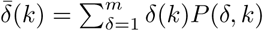. Here *P* (*δ, k*) is computed via the probability distribution derived from expression (6). The variation of 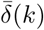 is displayed in Fig. S2. Notice how for small values of *k* the mean distance is close to *k*. For example, for *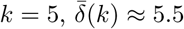*.

## RESULTS

Two distinct objectives are simultaneously addressed to characterise the metapopulation dynamics with facilitation under habitat destruction: (i) the impact of spatial correlations and distinct dispersal ranges (distances) of colonisation; and (ii) the effect of stochasticity in the metapopulation dynamics. However, before exploring these features, we will analyse the mean field model previously derived from the master equation. This simple model, which does not consider spatial correlations and is deterministic, will provide clues about the nature of the transitions involving metapopulations collapses under habitat degradation and pair-wise (facilitated) interactions among the individuals of the metapopulation.

As previously mentioned, two similar spatial models will be investigated. The first model considers facilitation during the process of plant reproduction and further colonisation, and the establishment of the seeds in the new colonised sites will not depend on the nearest neighbours at the colonising site. The second model will consider the process of facilitation necessary for the establishment of seeds in the colonised sites, being the reproduction of the plants not dependent on the density of the nearest neighbours.

### How facilitation determines metapopulation dynamics and extinction transitions in well-mixed populations under habitat destruction

The calculation of an explicit, closed solution *x*(*t*) is not possible for Eq. (5), as a difference from the classical Levins’ metapopulation model with habitat destruction (see Section S1). However, it is possible to qualitatively investigate the dynamics of the model incorporating facilitation. To do so, we will compute the equilibrium points and their stability, as well as a potential function. The equilibrium points of this model are obtained from *f* (*x*) = *cx*^2^(1 –*D– x*)–*ex* = 0, which gives three solutions, namely *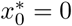* and the pair

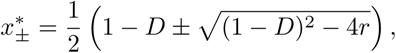

where *r ≡ e/c*. Notice that the pair *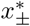*will be biologically meaningful whenever (1–*D*)^2^ –4*r ≥* 0. Indeed,when (1 - *D*)^2^ *-* 4*r* = 0 both equilibrium points *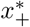* and 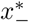 have the same value, which occurs for the critical threshold

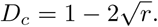

associated to a catastrophic shift, as shown in Fig. 2a. For *D > D*_*c*_ the term inside the square root of the equilibrium points *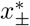* is negative (i.e., imaginary) and the two equilibria do not exist in the (biologically-meaningful) phase space of the real numbers, involving a catastrophic shift towards extinction occurs through a fold or saddle-node bifurcation (to be compared with the Levins’ model with habitat destruction, for which the extinction of the metapopulation is smooth and continuous, see Section S1).

**FIG. 2.**
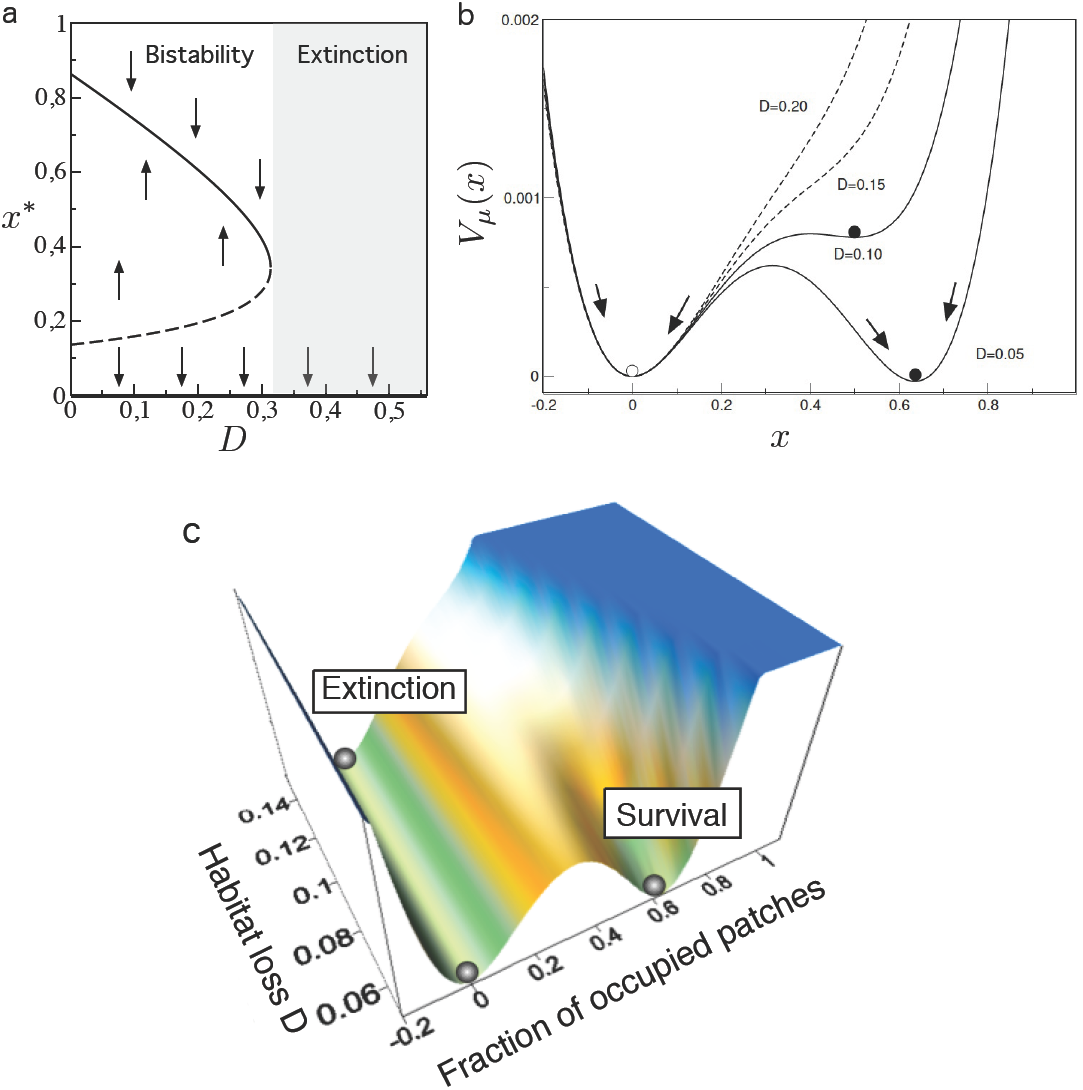
Catastrophic shift due to habitat destruction in metapopulations with facilitation identified in the mathematical model given by Eq. (5). (a) A tipping point is found as the fraction of habitat destroyed, *D*, increases beyond the bifurcation value *D*_*c*_ (here with 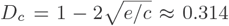 since *c* = 0.85 and *e* = 0.1) causing a saddle-node or fold bifurcation. (b) Another approach to this type of bifurcation diagram is provided by the potential function *V* (*x*) associated to the dynamics. Values of *D* below the bifurcation value (solid lines) display bistability, while for *D > D_c_* a single, stable state is found at *x*\* = 0. In (c) we show a more complete picture of the role played by *D* in changing the shape of *V* (*x*).

Generically, the (linear) stability of an equilibrium point *x*\* in a one-variable dynamical system can be obtained by means of the sign of *λ*(*x*\*) = *df* (*x*\*)*/dx*. The equilibrium point involving metapopulation extinction is a local attractor since *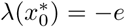*. This means that the stability of the origin does not depend on *D*, which plays the key role in the fold bifurcation described above. We refere the reader to Section S2 for further analyses on the stability of the equilibrium points *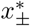*.

This model also allows for an explicit calculation of the associated landscape, described by the potential function *V* (*x*) = *−∫* (*x*)*dx*, which for Eq. (5) gives:

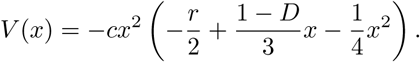

The potential is displayed in Fig. 2b for several values of *D*. For *D < D*_*c*_ (solid lines) two wells are found, corresponding to the two stable states (*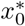*and *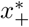*) resulting in bistability, that will be achieved depending on the initial conditions. Interestingly, evidences of multiple states have been recently identified in semi-arid ecosystems (Berdugo *et al.* 2017). Also, a complete fold bifurcation diagram has been experimentally built with cooperative yeast strains (Dai *et al.* 2012). Once the fraction of habitat destroyed surpasses its critical values *D*_*c*_, a single well is found (dashed lines). This single stable state is given by the equilibrium point *x*\* = 0, and involves the extinction of the metapopulation. A three-dimensional visualisation of the function *V* (*x*) is plotted for different values of *D* in Fig. 2c. Here, it is possible to see the dependence of the initial conditions (fraction of occupied sites) in the asymptotic dynamics under the bistable scenario (survival and extinction).

### Spatially-explicit dynamics and transitions to extinction with facilitation in reproduction under habitat destruction

In this section we will further extend the results obtained from the mean field model, which assumed well-mixed dynamics and determinism. Here we will use the cellular automata (CA) models to include explicit spatial correlations and stochasticity. In what follows we will denote the normalised population of individuals (occupied sites) at generation *τ* as *ρ*(*τ*). The term *ρ*\* will denote population at equilibrium (i.e., large *τ*). We first investigate the dynamics of facilitation during the process of reproduction and colonisation. This means that seeds can occupy non-destroyed empty sites, but their growth and further reproduction will be determined by facilitation provided by the presence of closer neighbours. This mode of facilitation involves that a given individual needs from other individuals in its surroundings to further reproduce and colonise. Such neighbouring individuals may retain water and provide good conditions for the growth and reproduction of the species after the arrival of seeds by dispersion. The dependence of neighbours for reproduction might also be representative of dioecious species such as mediterranean shrubs e.g., *Pistacia lentiscus* (Quezel 1981) or *Juniperus sp.* (Adams 2004), as well as plants of the *Salix* genus (Newsholme 1992) or African teaks e.g., *Baikiaea plurijuga, Milicia excelsa, Pericopsis elata*, or *Pterocarpus angolensis*, among others. For these cases, the reproduction and colonisation term will be density-dependent (non-linear), as modelled by Eq. (5), since it is assumed that plants will be fertilised by the surrounding ones (assuming short-range fertilisation).

We start analysing two extreme scenarios of colonisation: (i) the new individuals are dispersed to any random site of the lattice; (ii) the new individual colonises the nearest neighbour of the selected site (a neighbour at a Chebyshev distance of 1 i.e., Moore neighbourhood for colonisation with dispersal to the 8 nearest neighbours). Scenario (i) is closer to the mean field model analysed above, since here spatial correlations are only considered during the process of facilitation, but then colonisation can take place to any site of the lattice (similar to the break of the spatial correlations). For this type of colonisation a catastrophic shift is found, as predicted by the mean field model (see Fig. 2a). Figure 3 displays the mean population density *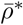*for different values of habitat destruction and colonisation probabilities. The system with long-range colonisation displays a catastrophic shift and bistability, as predicted by the mean field model. Figure 3a displays the abrupt change from survival to extinction, which takes place at increasing *D* and at decreasing *c*. The time series in Fig. 3a.1-a.2 shows the bistability of the system in the survival scenario: low initial conditions give place to metapopulations extinctions (Allee effect) while large initial populations allow the survival of the metapopulation (see also Figs. S3-S6).

**FIG. 3.**
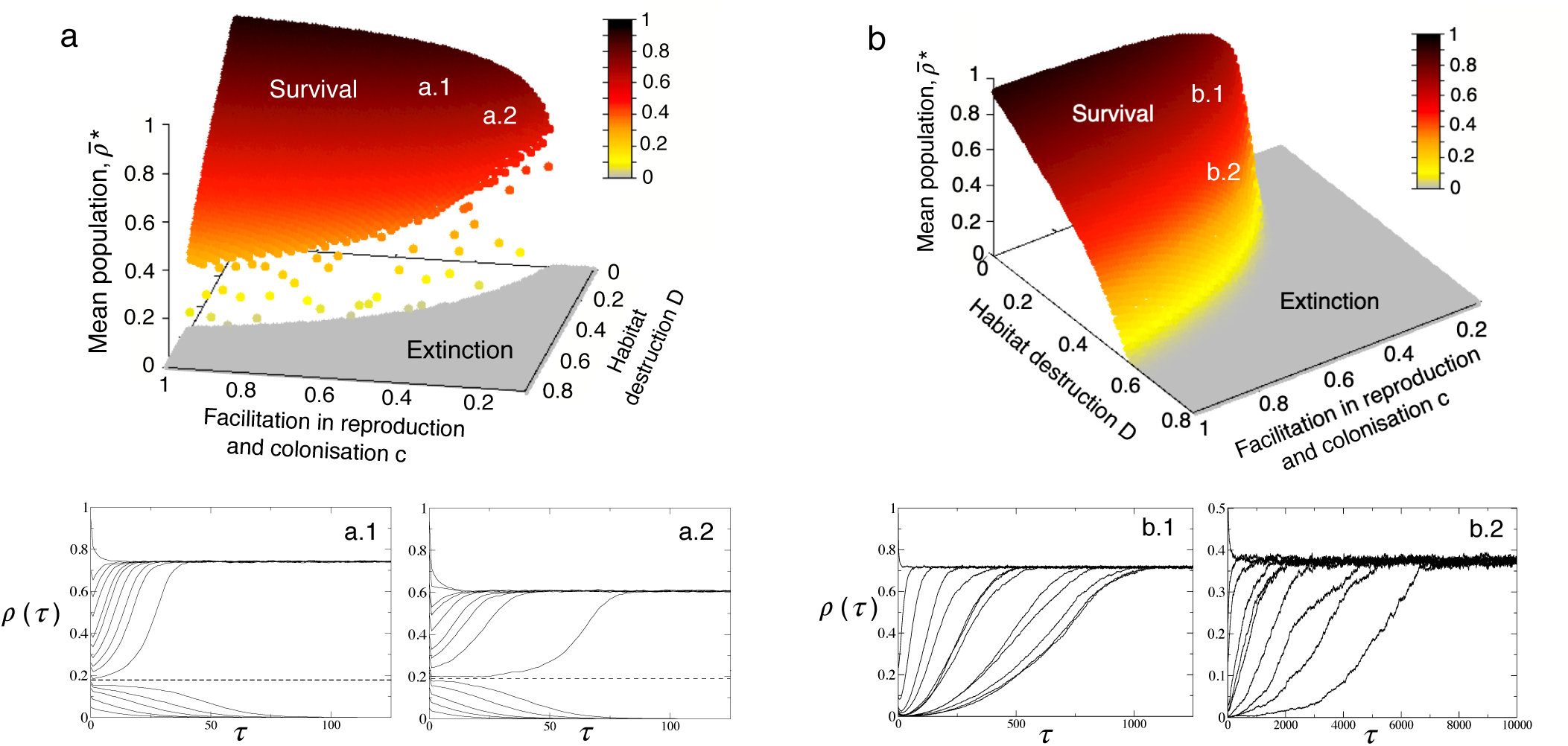
Transitions due to habitat destruction in the spatial model with facilitation in reproduction before colonisation for (a) long-range (using 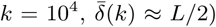 and (b) local dispersal (using 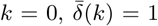), respectively. The surface are the; stationary population values for different values of *D* and *c* (here we use *e* = 0.05). Each data point is an average over 25 independent replicates at *τ* = 5 *×* 10^4^ using *L* = 100. Below we display two examples of the time dynamics for both long-range and local dispersal using a battery of different initial conditions, using the combination of parameters (*c, D*) placed on the surfaces with letters a and b in white. For long-range colonisation, the dynamics depends on the initial conditions, since for the same value of *c* and *D*, survival or extinction can be achieved due to bistability, in agreement with the mean field model (panels a.1 and a.2). For the local dispersal, however, the system becomes monostable and undergoes a continuous transition (panels b.1 and b.2).

Surprisingly, the same analysis allowing only for local colonisation *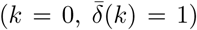* reveals a continuous transition towards extinction (Fig. 3b). For this case, bistability is not found, since all initial conditions in the survival scenario allow the persistence of the metapopulation (see the time series in Fig. 3b.1-b.2). Our results suggest that the dispersal distance is crucial in metapopulations with facilitation, since it can determine if transitions due to e.g., habitat destruction are catastrophic or continuous. An example of the abrupt transition occurring when dispersal is completely random and the associated spatio-temporal patterns are displayed in Fig. 4. Here, we plot the mean population value,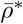, at increasing the fraction of habitat destruction, *D*, setting *L* = 200 (Figs. S3a displays the same analysis using lattices of sizes *L* = 50, 100, and 150). For low values of *D*, the metapopulation is able to persist, and the observed spatial patterns are well-mixed. However, even for low *D* values, extinction can occur if the initial population values are low (see Figs. S3a, S4a and S6). This is due to the existence of a separatrix corresponding the unstable branch identified with the mean field model (given by the repulsor point *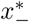*). Section S3.2.1 explains how this separatrix can be systematically computed in the CA model (see below). Once the critical value of *D*_*c*_ is surpassed, the metapopulation suffers the catastrophic shift. Notice that near *D*_*c*_ the metapopulation suffers a long transient (with plateau shape, as can be seen in the mid time series of Fig. 4) before collapsing. This is a fingerprint of saddle-node or fold bifurcations, where delayed transitions are known to occur near bifurcation thresholds (Sardanyé s & Solé 2007, Fontich & Sardanyé s 2008). Once *D* grows beyond *D*_*c*_ the transient to extinction becomes very short.

**FIG. 4.**
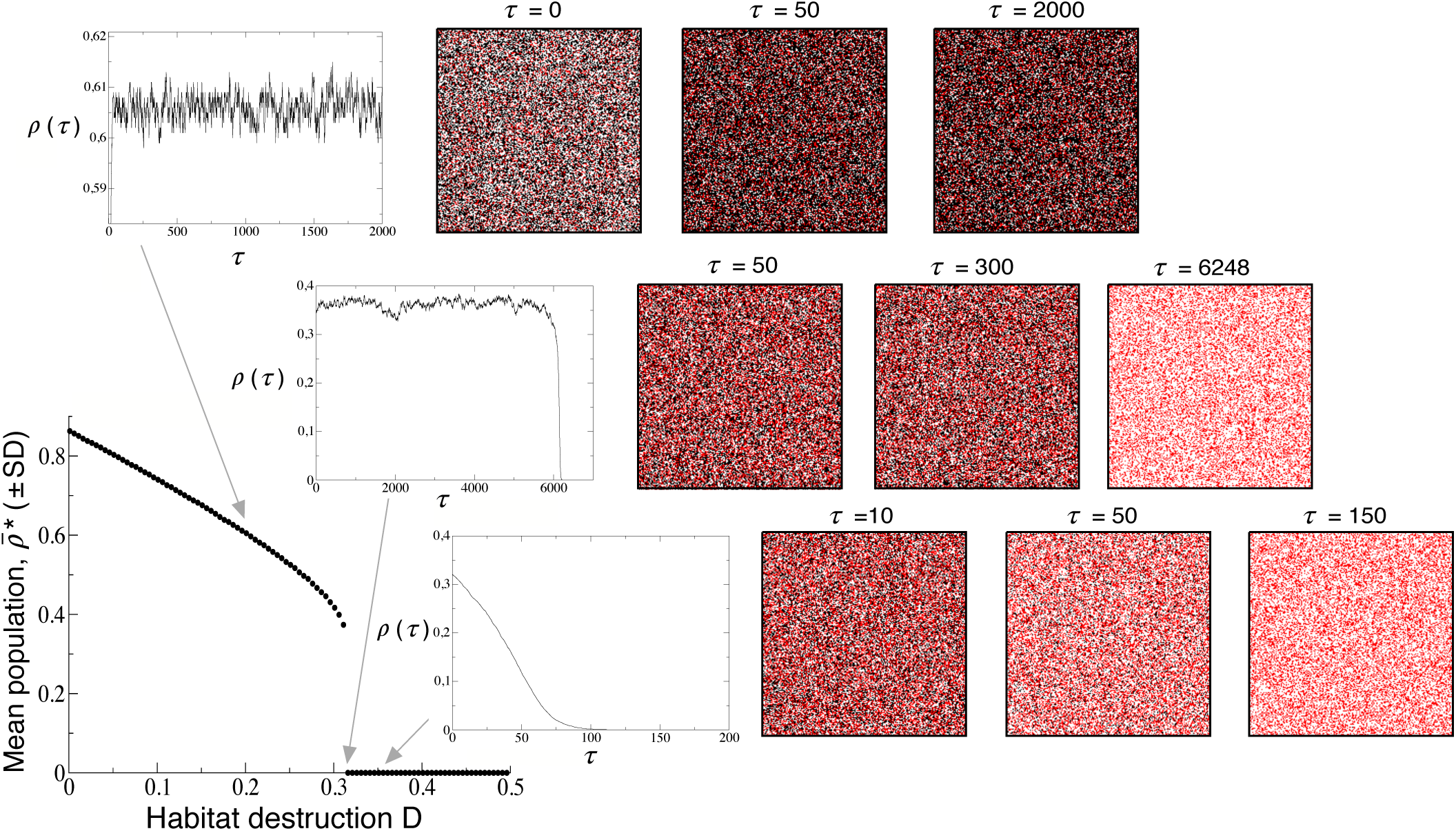
Spatio-temporal dynamics of facilitation considering random colonisation (using *k* = 10^4^ i.e., any lattice site can be colonised with equal probability) setting *c* = 0.85, *e* = 0.1, and *L* = 200 (see Fig. S3a for the same analysis using smaller lattice sizes). The mean equilibrium population values (*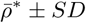*, error bars are smaller than the points) are computed averaging 25 independent runs at *τ* = 5 ×10^4^. We display three examples of different fractions of habitat destruction, *D*, with (from top to bottom): *D* = 0.2, *D* = 0.3122 ≈ *D_c_*, and *D* = 0.35. The spatial patterns display several snapshots at different generations of the CA. Here we represent the sites occupied by an individual (black), the destroyed sites (red), and the empty sites (white).

The metapopulations with local colonisation form patchy aggregations, which become more disconnected and dispersed as *D* approaches to the critical threshold. Figure 5 displays the same results of Fig. 4 but now simulating local colonisation. Here, although extinction occurs for a similar *D*_*c*_ value compared to the long-range colonisation analysis, the transition is smooth (see also Fig. S4). Similar analysis using lattices with *L* = 50, 100, and 150 are displayed in Fig. S3b. The algorithm used to find the separatrix in the CA have been also applied for local colonisation. Here, the separatrix has not been found, suggesting that the system under local colonisation becomes monostable.

**FIG. 5.**
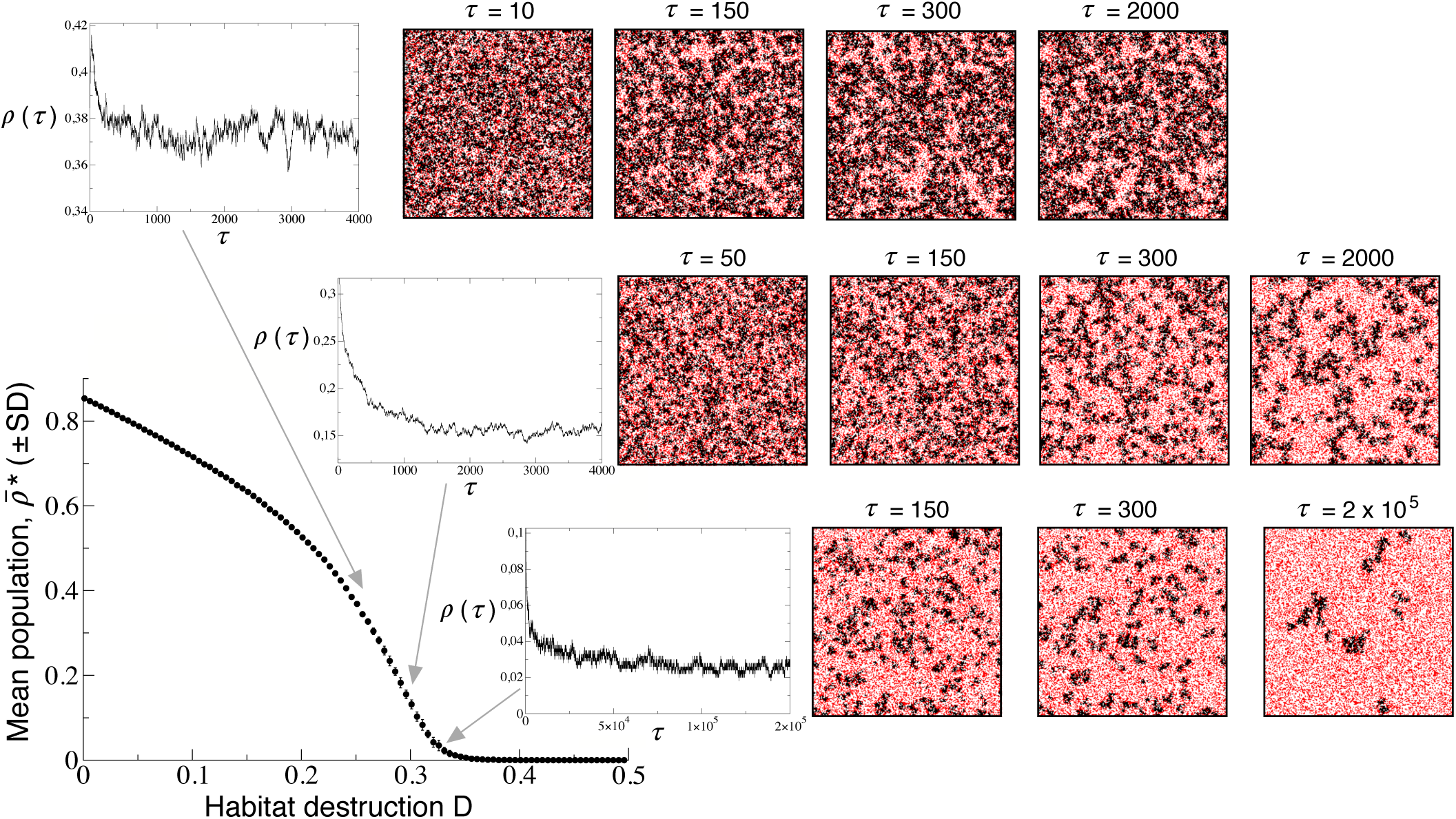
Same as in Fig. 4 now considering facilitation and local colonisation within the nearest neighbours (setting *k* = 0). Here we also used *c* = 0.85, *e* = 0.1, and *L* = 200 (see Fig. S3b for the same analysis using smaller lattice sizes). We also display several examples of the spatio-temporal dynamics for four different fractions of habitat destruction, *D*, with (from top to bottom): *D* = 0.25, *D* = 0.3, and *D* = 0.33. At incresing *D*, patchiness appear and become more sparse.

### How does the range of seed dispersal influence the extinction transitions under facilitation in reproduction under habitat destruction

The previous results have focused on two extreme cases of colonisation given by purely random (long-range) colonisation and local dispersal (involving the colonisation of the nearest neighbours). However, several interesting questions arise from the previous results: how does the nature of the transition to extinction depend on the range of dispersal? What is the impact of the extinction probability (*e*) on the nature of the transitions for different ranges of dispersal? To address these questions we will simulate the spatial dynamics setting *k >* 0 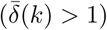. Figure 6a displays results on the mean population values increasing the fraction of destroyed habitat, *D* (notice that we focus on a range of *D* values close to the extinction threshold). As mentioned, he results for *k* = 0 the transition is smooth and the critical *D* value is about *D*_*c*_ *≈* 0.35 (Fig. 6a), since the density of individuals decreases in a flat manner. However, when *k* = 6 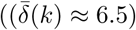 the transition becomes more abrupt, being *D*_*c*_ ≈ 0.32. The inset in Fig. 6a displays the same results for 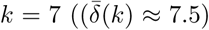 and 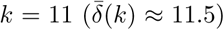, where more abrupt transitions are found. Figure S5a displays the same results for *k* = 0, *…*, 11. The previous results also indicate that the critical value *D*_*c*_, which is tied to transitions of different nature, can change for different distances of colonisation (see Fig. 6a and 6b, and Figs. S5a and S6). Although the differences are not very large, increasing the range of dispersal decreases *D*_*c*_. However, this is not a general trend, since increasing *e* can reverse this situation (compare Fig. 6a with Fig. 6b). This effect is discussed in more detail below.

**FIG. 6.**
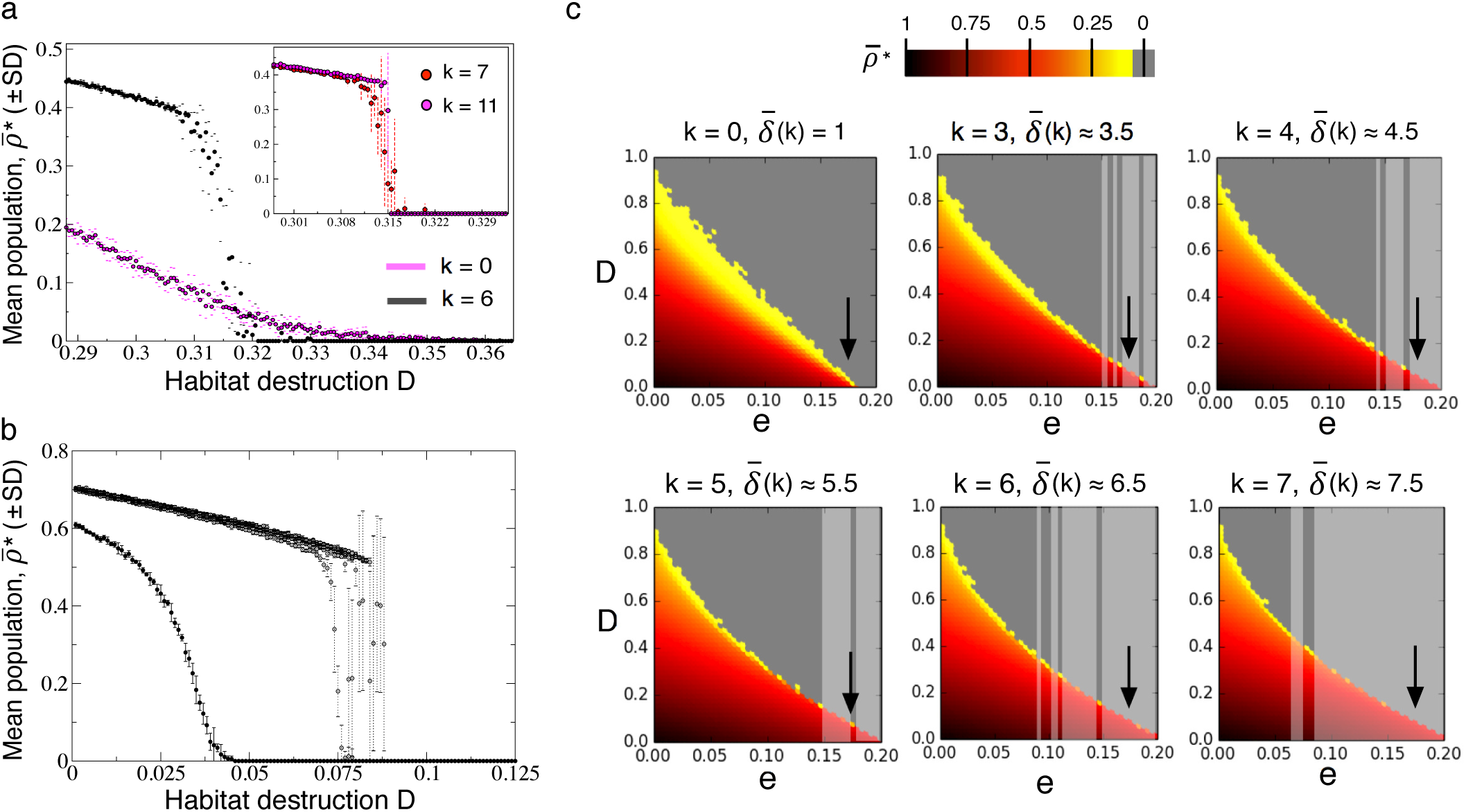
(a) Transitions towards extinction at increasing the fraction of destroyed sites using several colonisation ranges changing *k*, with *c* = 0.85 and *e* = 0.1. Here each data point is the mean population 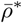 (and the small lines are the extremes of the ± *SD*) averaged over 5 independent replicas at τ = 2 ×10^5^. The main panel displays the values for *k* = 0 (local colonisation) and *k* = 6, while the inset displays the same results for *k* = 7 and *k* = 11 (here the *SD* is displayed with dashed lines). Notice that for this combination of *c* and *e* probabilities, the continuous transition becomes discontinuous *k* ≈ 6 (see Fig. S5(b) for the same analyses using *k* = 0, *…*, 11). (b) Same as in (a) using *e* = 0.175 and different colonisation ranges (from left to right): *k* = 0 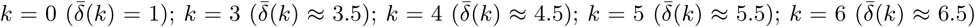; and *k* = 7 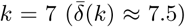. (c) Mean population values in the parameter space (*e, D*) using *c* = 0.85 and six different values of *k* (*k* = 0, 3, 4, 5, 6, 7). Here each data point is also the average of 5 independent runs now setting τ = 2 × 10^4^. The overlapped grey bands enclose the regions where abrupt transitions are found. Notice that the change from yellow to gray corresponds to the continuous transition, while the change from red (orange) to grey accurately displays the sharp transition. In all of the plots we used *L* = 200.

In order to distinguish between continuous and abrupt transitions several strategies could be followed. One possible approach, as previously mentioned, could be the computational search of the so-called separatrix, which is given by the unstable branch in the bistability scenario before the saddle-node bifurcation (see dashed line in Fig. 2a, and Figs. S3a and S4). The separatrix divides the basins of attraction of the extinction and the persistence attractors. We have developed an algorithm for finding this separatrix in the CA model. The description of this method can be found in Section S3.2.1. For the probabilities of colonisation and extinction (*c* = 0.85 and *e* = 0.1) analysed, the change from continuous to catastrophic transition is shown to take place at *k* = 6 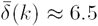(see Fig. S6 and Section S3.2.1 for the explanation of these results).

Up to now, we have determined that the nature of the transitions can change depending on the distance of dispersal. However, as we have previously seen, these transitions can also depend upon the model parameters (i.e., colonisation or extinction probabilities). In order to explore the impact of the model probabilities we will focus on the impact that extinction probability *e* has in the critical *D*_*c*_ values and in the nature of the transitions. Figure 6b displays the mean population values also tuning *D* for different values of *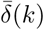*, using a larger extinction probability *e* = 0.175. For local colonisation (with 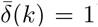) the transition is continuous and metapopulation extinction occurs at *D*_*c*_ *≈* 0.04. However, the increase in the dispersal distance changes the continuous transition towards a more abrupt one, and the value of *D*_*c*_ increases up to *D*_*c*_ ≈ 0.08 (Fig. 6b). This counterintuitive result highlights the importance of the difficulties in predicting the changes of the system under nonlinearities together with spatial and stochastic dynamics.

Figure 6c displays the impact of the dispersal distance and extinction probabilities (analyses in the parameters space (*e, D*)) in the mean density of the metapopulation as well as in the nature of the transitions. For local colonisation, the transition is shown to be continuous for all the analysed ranges of *e*, here with 0≤ *e≤* 0.2. Notice that a continuous transition occurs in the boundaries of the triangle inside the space (*e, D*) with yellow colour, since yellow colour indicates close to zero values and thus no catastrophic jump is found. For all of the boundaries with orange or red colour the jump can be considered abrupt. Specifically, those values of *e* causing a catastrophic transition has been framed with a transparent grey area in Fig. 6c. The arrows in the panels of Fig. 6c indicate the transitions for *e* = 0.175, which are displayed in Fig. 6b plotted with standard deviation indicated with dashed lines. Finally, in Fig. S5b we display the change in the mean population values at increasing the extinction probability considering local colonisation. Here we must notice that the the catastrophic shift only takes place for *D* = 0. Once the habitat starts to be destroyed the transition switches to a continuous one.

### Spatially-explicit dynamics and extinction transitions with facilitation in seeds establishment under habitat destruction

A second model considering facilitation during the establishment after colonisation is analysed here. In the previous model, seeds were able to colonise non-destroyed, empty sites and then reproduce in a density-dependent manner. In the system investigated here, facilitation will enhance the establishment of the seeds, finding good conditions for its establishment. However, reproduction will not depend on facilitation (see Fig. S7 for a schematic diagram of this system). As we will show below, the dynamics for this system is similar to the one previously analysed, although some differences in the extinction transitions are found.

Now the CA model needs to be slightly modified to simulate these ecological processes. The state-transition rules of this model are explained in Section S4.1. Although similar to the previous model, facilitated establishment becomes more sensitive to extinction. For instance, the impact of extinction rates, *e*, in the mean population is stronger compared to the system previously analysed. In Fig. S8 we plot the mean population of occupied sites increasing extinction probabilities using different fractions of habitat destruction. The extinction thresholds are found for lower values of *e* and the same tendencies are observed for decreasing values of *D*. We notice that transitions for *D >* 0 are also continuous, while the extinction for *D* = 0 is abrupt, as we found in the previous model with facilitation in reproduction. The value at which the catastrophic shift takes place for this second model is *e* ≈ 0.164, compared to the value obtained for the model with facilitated reproduction given by *e ≈* 0.184.

The differences in the critical values of the probabilities causing metapopulation extinction for this second model are also observed tuning the fraction *D* of habitat destruction. These differences can be seen in Figs. S9 and S10, where we plot the mean population of occupied sites increasing *D*. These data are shown overlapped to the same analysis obtained with the model considering facilitation in reproduction (displayed with small grey circles). Figure S9 displays the results for long-range colonisation. For this particular analysis, the extinction occurs at *D ≈* 0.26 for the model with facilitated establishment, while the critical *D* value for the model with facilitated reproduction is *D ≈* 0.315. The spatial patterns for long-range colonisation are also random like distributions of the occupied sites (Fig. S9). This second model with local colonisation also displays patches of occupied sites, although the distribution of these patches is more sparse compared to the same analyses developed for the model with facilitated reproduction (Fig. 5) due to the higher sensitivity of this second system (see Fig. S10).

## DISCUSSION

Local facilitation among individuals in a spatially-extended landscape pervades the presence of strong nonlinearities and the potential for tipping points. Because of the underlying higher-order interactions, non-monotonous responses to changing environmental conditions are expected to occur. One particularly relevant case study is provided by semiarid ecosystems. As grazing or aridity increase, vegetation and soil quality levels decay until breakpoints are reached, resulting in catastrophic shifts (Kéfi *et al.* 2007a-b). On the other hand, habitat los and fragmentation interacts with other types of nonlinearities associated to the presence of facilitation effects. In this paper we brought together both habitat loss and positive interactions with the goal of exploring the potential transitions to extinction. Both well-mixed (non-spatial, mean field) and spatially extended versions are considered.

Our study considers both analytic and simulation model approaches. A derivation of the minimal (deterministic) model incorporating both positive interactions and fragmentation has been derived, leading as a limit case our generalised Levins model. This model allows to understand the expected impact of both habitat loss (as given by the *D* parameter) and facilitated colonisation (given by *e*). A closed analytical result is provided that gives a potential function predicting a sudden shift once a habitat destruction threshold is reached. The interesting first result here is that the threshold is different from the one predicted from the standard model incorporating habitat loss and competition. In the later the transition is continuous, whereas in our model it leads to a discontinuous one due to a fold bifurcation.

By extending the model to a spatial, stochastic scenario, a very interesting phenomenon has been found. When long-range dispersal is considered, a discontinuous shift is still found. Departures from the mean field predictions are known to occur when the role played by space becomes explicit. However, in this case the difference goes far beyond a displacement of the parameters associated to the transitions: the nature of the transition itself is changed and no shift is observed separating the survival from the extinction phase in the (*D, c*) parameter space. Such qualitative change in the nature of the transition resulting from short versus long dispersal is likely to be similar to those found in the study of other nonlinear systems under dimensionality changes. This has been studied in models of trimolecular reactions (Provata *et al.* 1993; Prakash & Nicolis 1996, 1997; Windus and Jensen 2007). More recently, a theoretical analysis of the so called Ginzburg-Landau model (Weissmann and Shnerb, 2014):

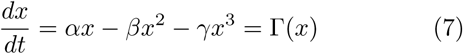

have been studied in order to explore catastrophic desertification (Weismann *et al.* 2017) as well as potential ways of avoiding catastrophic shifts (Vila Martin *et al.* 2018). Clearly our model belongs to this general class, mapping our parameters into the previous ones as follows:

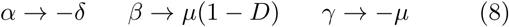

By expanding this model into a stochastic spatial counterpart, it is possible to actually explore the effect of different factors on the nature of transitions (Goldenfeld 2018). This was done by analysing the behaviour of

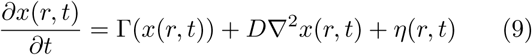

where the two last terms at the right hand side of Eq. (9) are the diffusion and noise terms associated to the spatiotemporal dynamics. Here *r* indicates a spatial coordinate. The paper predicts that limited diffusion (which they can treat as a continuous parameter) can transform the shift into a continuous phase change, as it occurs in our model. The analysis in Vila Martin *et al.* (2015) is obtained by means of the renormalisation group, a powerful method that is using in statistical physics to identify universal phenomena in spatially distributed systems.

In their analyses these authors looked at factors influencing a suppression of catastrophic shifts. In our study, instead, we can actually look at this in an adaptive context. The risks associated to vegetation loss due to grazing or aridity (factors increasing soil degradation) might have pushed species living in semiarid contexts to evolve local dispersal that will lead to patchiness while effectively removing the presence of a breakpoint. A smooth transition would not only prevent severe population extinctions but also favour, on a shorter and local scale, the persistence of vegetation cover.

## Acknowledgments

We would like to thank the members of the Complex Systems Lab as well as Maestre’s Lab from Universidad Rey Juan Carlos for very stimulating and useful discussions. We also want to acknowledge Antoni Guillamon, Ernest Fontich, and Jorge Duarte for helpful suggestions. This researchk has been supported by the Botìn Foundation by Banco Santander through its Santander Universities Global Division, a MINECO FIS2015-67616 fellowship and by the Santa Fe Institute. This work has also counted with the support of Secretaria d’Universitats i Recerca del Departament d’Economia i Coneixement de la Generalitat de Catalunya. The research leading to these results has received funding from “la Caixa” Foundation. This work has been also partially funded by the “Marìa de Maeztu” Programme for Units of Excellence in R&D (MDM-2014-0445), and from the CERCA Programme of the Generalitat de Catalunya. JS has been also funded by a “RamÓn y Cajal” Fellowship (RYC-2017-22243).

